# rhinotypeR: An R package for Rhinovirus Genotyping

**DOI:** 10.1101/2025.07.14.664540

**Authors:** Martha M. Luka, Ruth Nanjala, Wafaa M. Rashed, Winfred Gatua, Olaitan I. Awe

## Abstract

Rhinoviruses (RV) are common pathogens characterized by extremely high antigenic and genotypic diversity, yet the tools for their genotyping remain limited. We present rhinotypeR, a Bioconductor package designed to streamline the genotyping of RVs using the VP4/2 region by automating sequence comparison against prototype strains and applying predefined pairwise distance thresholds. RhinotypeR offers a comprehensive suite of functions, including sequence alignment, genetic distance calculation, and genotype assignment, which collectively simplify the often convoluted process of RV classification. Additionally, the package supports visualization tools for single-nucleotide polymorphisms (SNPs), amino acid substitutions, and phylogenetic relationships, providing an accessible platform for genetic analysis and evolutionary studies of RV. Comparative analyses demonstrate that genetic distance calculations by rhinotypeR align closely with established software such as MegaX and ape. RhinotypeR addresses the need for accessible and robust genotyping tools by offering a suite of functions that support the workflow of rhinovirus genetic analysis. It empowers researchers with a toolkit that enhances data processing efficiency and provides detailed, actionable insights into the genetic diversity and evolution of rhinoviruses.

## 1. Introduction

Genomic studies have transformed our understanding of viral infections, providing invaluable insights into their transmission pathways and evolutionary dynamics^1–5^. With the surge of sequence data availability, the capacity to dissect the complex mechanisms underlying virus spread and mutational landscapes has become increasingly important^6^. This is particularly key for pandemic preparedness - a matter of ‘when’ rather than ‘if’-highlighting the need for analytical tools that enable rapid responses to emerging infectious threats, from data acquisition to actionable insights^7–10^, ultimately contributing to global health security.

Rhinoviruses (RV) stand out as one of the most prevalent human respiratory pathogens^11–13^, yet their impact on human health is often underestimated^14,15^. Their genome is roughly 7.2 kb and consists of a single open reading frame that encodes for 11 proteins: seven nonstructural proteins (2A, 2B, 2C, 3A, 3B, 3Cpro and 3Dpol) and four structural proteins (VP1, VP2, VP3, and VP4)^15^. RV are classified into three species (RV-A, -B, and -C)^16^, which are further classified into an expanding tally of genotypes, currently totalling 169^17^. These genotypes exhibit unique antigenic properties^18^ and undergo independent evolutionary paths^19^, posing challenges for tracking RV transmission and evolution. The current classification of RV into genotypes relies on the analysis of the VP1 or VP4/2 genomic regions, which have demonstrated congruence in reflecting RV genetic diversity^20,21^. However, the VP4/2 is more commonly used due to its shorter length and less variability, hence ease of amplification^21^. Sequences are typically aligned with prototype strains ^17^, and classification is based on pairwise distances to these prototypes, adhering to specific thresholds for each RV species (<10.5% for species A and C, <9.5 for species B)^16,20,21^. Although this approach sounds simplistic and straightforward, it can be a convoluted process even for the same dataset that is processed by different individuals.

Despite the abundance and burden of rhinoviruses, we lack a streamlined approach to genotype infections using the VP4/2 region. We therefore introduce rhinotypeR (https://bioconductor.org/packages/release/bioc/html/rhinotypeR.html), a Bioconductor package that addresses this challenge by providing a platform for the rapid and accurate classification of RV to the genotype level. By automating the comparison of sequence data against prototype strains and implementing predefined pairwise distance thresholds, rhinotypeR streamlines the genotype assignment process. This package enhances our epidemiological toolkit, enabling more efficient surveillance and analysis of rhinoviruses.

## 2. Methods

### 2.1. Workflow

The rhinotypeR package offers a detailed and streamlined workflow for rhinovirus genotyping and analysis, as illustrated in **Figure 1**. Starting with the retrieval of RV prototype sequences using *getPrototypeSeqs()*, the package allows the user to download RV prototypes in a FASTA format into a local directory, ensuring easy access for subsequent analysis. GenBank accessions of the prototype sequences used are listed in **Supplementary Table 1**. To accurately assign RV types, the user should combine the downloaded prototypes with their newly generated sequences before alignment. This will help ensure that the input reflects the correct genomic section of the VP4/2 sequences. For functions other than *assignTypes()*, the input sequences do not need to contain the prototypes. However, a high-quality alignment remains critical.

**Figure 1.**
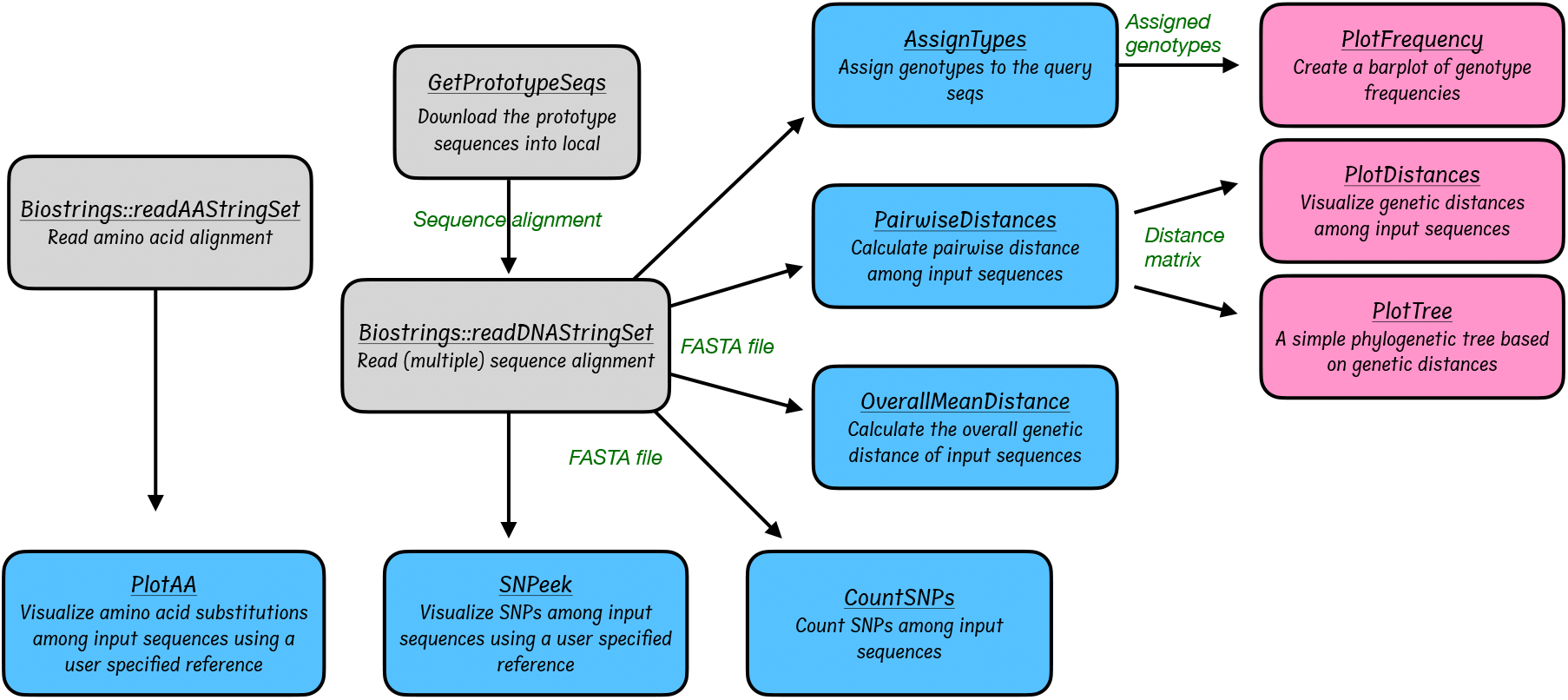
Workflow of the rhinotypeR package. The process begins with the user downloading prototype sequences included within the package. These are then aligned with newly generated sequences to create a comprehensive dataset. The combined dataset is uploaded into R for downstream analysis and visualization.

#### Data

The package takes in RV sequences in FASTA format. While several alignment tools are proficient, we recommend using the default settings of MAFFT software^22^ for alignment. This preference is based on our positive experiences with MAFFT’s speed and accuracy in aligning virus sequences. Following the initial alignment, we advise manually curating the alignment to truncate any genomic regions outside the required VP4/2 region and realign as necessary. Alternatively, one can also include arguments in the alignment software to remove bases before the first and after the last base of a reference sequence, where applicable. It is important to note that a suboptimal alignment can significantly distort the inference of mutations, which can falsely elevate or decrease the estimated genetic distances. Therefore, careful attention to alignment quality is crucial for robust genetic analysis.

#### Ensuring high-quality data input and accurate analysis

To maximize the accuracy and reliability of the rhinotypeR package analyses, it is essential to maintain high standards of data quality. Here are key considerations:

1. **Verification of sequence quality**. The user should assess the quality of each sequence during consensus calling. Remove or correct sequences with poor quality scores to prevent the introduction of errors into the analysis.
2. **Appropriate sequence lengths**. We recommend that sequence lengths be no shorter than 350 bases. The optimal length for VP4/2 sequences is approximately 420 bases, comprising about∼207 bases from VP4 and ∼213 bases from VP2.
3. **High-quality alignment**. We recommend that the user include the downloaded prototype sequences in their alignment to serve as references. Use robust alignment software or algorithms and check alignments to ensure accuracy.
4. **Minimize gaps**. Gaps, representing insertions and deletions, should be minimal in the sequence data. Excessive or incorrectly placed gaps can lead to significant errors in distance calculations and may misrepresent the evolutionary relationships among sequences.

#### Analysis

Once the prototypes are downloaded and combined with the query sequences, they are aligned and manually checked for accuracy. Users can subsequently import the curated VP4/2 alignment from a FASTA file directly into R using the *readDNAStringSet()* function from the Biostrings package ^23^, enabling downstream genetic analysis. The key functions of rhinotypeR and their required inputs and expected outputs are outlined in **Table 1**.

**Table 1.** RhinotypeR functions, their description, requirements and outputs.

| Function Name | Description | Input | Output format |
| --- | --- | --- | --- |
| *getPrototypeSeqs()* | Downloads RV prototypes into a user-specified local directory. | Destination path | RV prototypes are downloaded into the local machine |
| *SNPeek()* | Visualise single nucleotide polymorphisms (SNPs) using a predicted or user-specified sequence as the reference. | FASTA file | A plot highlighting SNPs per sequence |
| *plotAA()* | Visualise amino acid substitutions using a predicted or user-specified sequence as the reference | Amino acid FASTA file | A plot highlighting amino acid substitutions per sequence |
| *assignTypes()* | Genotype assignment to query sequence | FASTA file | CSV file with three columns: sequence header, assigned type, and genetic distance |
| *pairwiseDistances()* | Calculate pairwise distance among input sequences using a user-specified evolutionary model. | FASTA file | A dense distance matrix |
| *overallMeanDistance* | Calculates the overall mean genetic distance of query sequences using a user-specified evolutionary model. | FASTA file | A single numeric value |
| *countSNPs()* | Count pairwise SNPs among query sequences. | FASTA file | A dense matrix |
| *plotFrequency()* | Creates a barplot of genotype frequencies. | Assigned types | Barplot |
| *plotDistances()* | Plots prototype distances | Distance matrix | A heatmap |
| *plotTree()* | Plots a simple phylogenetic tree from the distance matrix. | Distance matrix | A simple phylogenetic tree |

The *pairwiseDistances()* function calculates genetic distances between all pairs of sequences using a user-specified evolutionary model, generating a dense distance matrix. This matrix is versatile, supporting analyses such as phylogenetic inference and multi-dimensional scaling. To summarize genetic diversity across sequences, users can employ *overallMeanDistance()*, which calculates a single numeric value representing the mean genetic distance among the sequences.

For genotype assignment, *assignTypes()* computes genetic distances and matches each query sequence to the closest prototype strain, provided the distance falls within predefined thresholds. The output includes four columns: the query sequence identifier, the assigned genotype, the genetic distance, and the corresponding prototype strain. Sequences that exceed the threshold distance from any prototype are marked as unassigned, indicating potential novelty, high divergence, or data quality issues, warranting further investigation.

#### Evolutionary models

The *countSNPs()* function counts pairwise SNPs among the query sequences, to provide a dense matrix that serves as a basis for estimating evolution. Genetic distances are calculated using any of the four models: pairwise distance (devoid of evolutionary assumptions), Jukes-Cantor (JC)^24^, Kimura 2 parameter (K2P)^25^, or Tamura 3 parameter (T3P)^26^ evolutionary models. Each model has its set of assumptions and limitations^27^, and the user should choose the best model based on their data.

The pairwise distance method is the simplest model, but it does not explicitly account for evolutionary processes. It assumes that genetic distances accumulate linearly over time ^28^. The Jukes-Cantor model, on the other hand, provides a simplified foundational approach to understanding evolutionary dynamics by assuming neutral evolution^24^. Meanwhile, the K2P and T3P offer more advanced methods to understand molecular evolution by incorporating differential rates for transitions and transversions, thus providing more realistic estimates of evolutionary distances than simpler models^25,26^.

While simpler models are faster, more complex models offer greater accuracy but require more data and computational effort. The best model isn’t always the most complex; it should fit the data and evolutionary questions^27,28^.

#### Visualization

RhinotypeR offers multiple visualization functions to aid the interpretation of genetic data. The *SNPeek()* function visualizes single-nucleotide polymorphisms (SNPs) among the input sequences, while *plotAA()* function visualizes amino acid substitutions, representing protein changes against a reference. An amino acid sequence alignment is read into R using Biostring’s^23^ *readAAStringSet()* function. For *SNPeek()* and *plotAA()*, the user specifies the reference sequence by moving it to the bottom of the alignment before importing it into R. The *plotFrequency()* function creates a barplot of assigned genotype frequencies, assisting researchers in understanding the frequency of each genotype, while *plotDistances()* visualizes prototype distances through a heatmap to compare genetic relatedness. Lastly, *plotTree()* constructs and visualizes phylogenetic trees based on the computed genetic distances, allowing researchers to interpret evolutionary relationships visually.

Our tool streamlines the genotyping process and integrates these analyses seamlessly into a coherent workflow, ensuring researchers can quickly and efficiently move from data acquisition to in-depth genetic analysis and visualization. This is crucial for advancing our understanding of rhinovirus genetic diversity and aiding in the global surveillance of RV infections.

### 2.2. Application to sample data and comparison to existing tools

We used 253 previously generated ^29^ VP4/2 FASTA sequences (GenBank Accessions MT177659– MT177911) to test the fidelity of rhinotypeR. These sequences represent a typical VP4/2 partial-genome dataset used to study rhinovirus dynamics. The sequences were of length 350 - 420 bases and were previously classified into 47 RV genotypes: 24 RV-As, 7 RV-Bs, 16 RV-Cs and two unassigned types ^29^. The sample data was processed with rhinotypeR to test the functions in **Table**

**1**. Genetic distances estimated by rhinotypeR were compared to commonly used software (MegaX

^30^ and ape ^31^) to assess fidelity and accuracy.

## 3. Results

### Overview, functionality and output

We present rhinotypeR^32^, a comprehensive tool aimed at streamlining the genotyping of rhinoviruses using the commonly used VP4/2 region and offers robust analytical and visualization functionalities. **Figure 2** displays outputs from key functions when applied to a typical rhinovirus VP4/2 dataset. Its primary function, *assignTypes()*, rapidly assigns genotypes and gives a simple, clear output that could guide further investigations.

**Figure 2.**
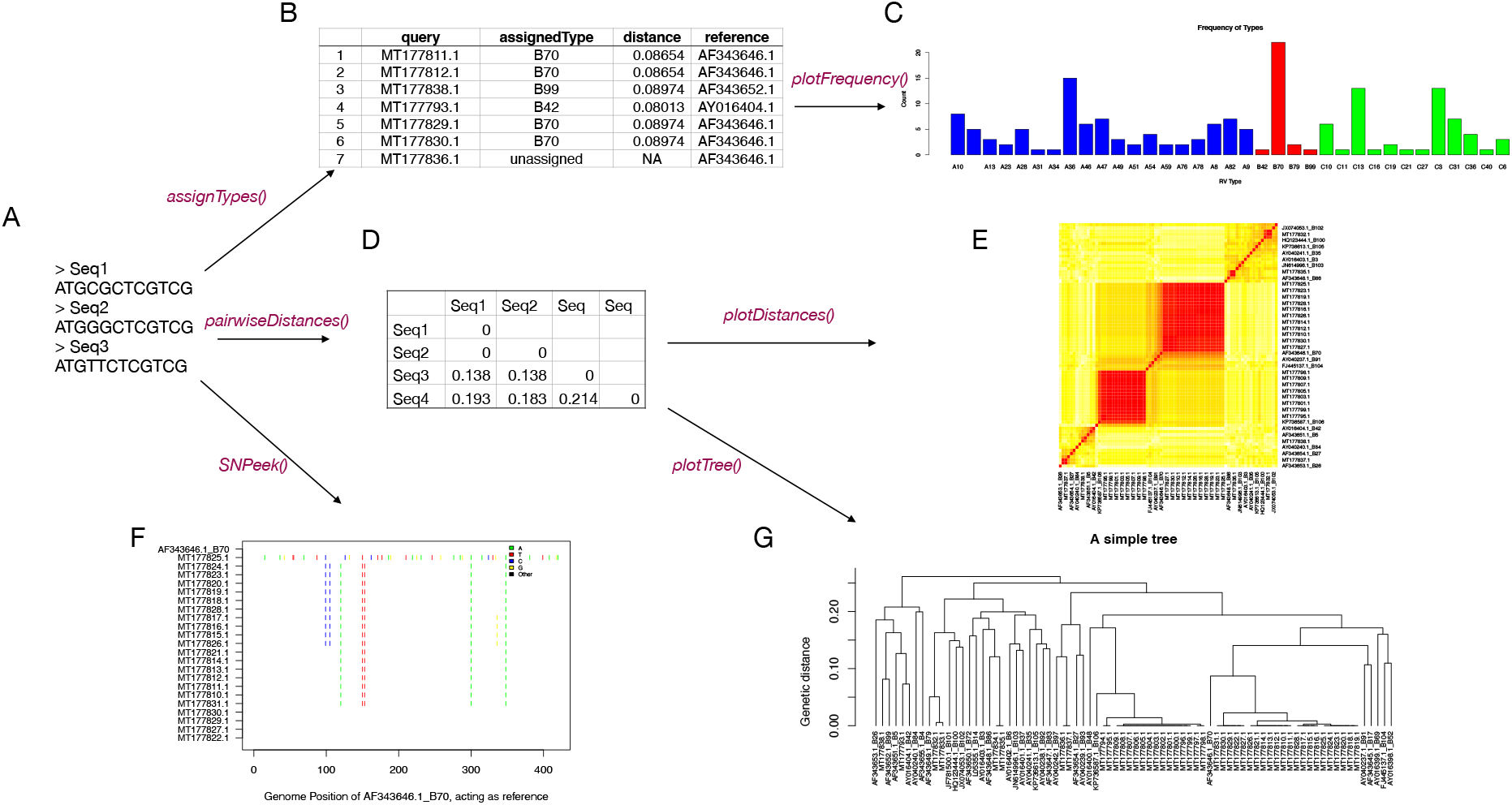
Key rhinotypeR functions and their outputs applied to a typical rhinovirus dataset. A) The user provides a curated alignment as input. The package can B) assign sequences to genotypes and C) plot the frequencies. D) The generated pairwise distances matrix can also be visualized as E) a heatmap. F) The user can visualize single nucleotide polymorphisms relative to a predicted reference. G) Simple phylogenetic analysis can also be performed to understand evolutionary relationships.

**Figure 3.**
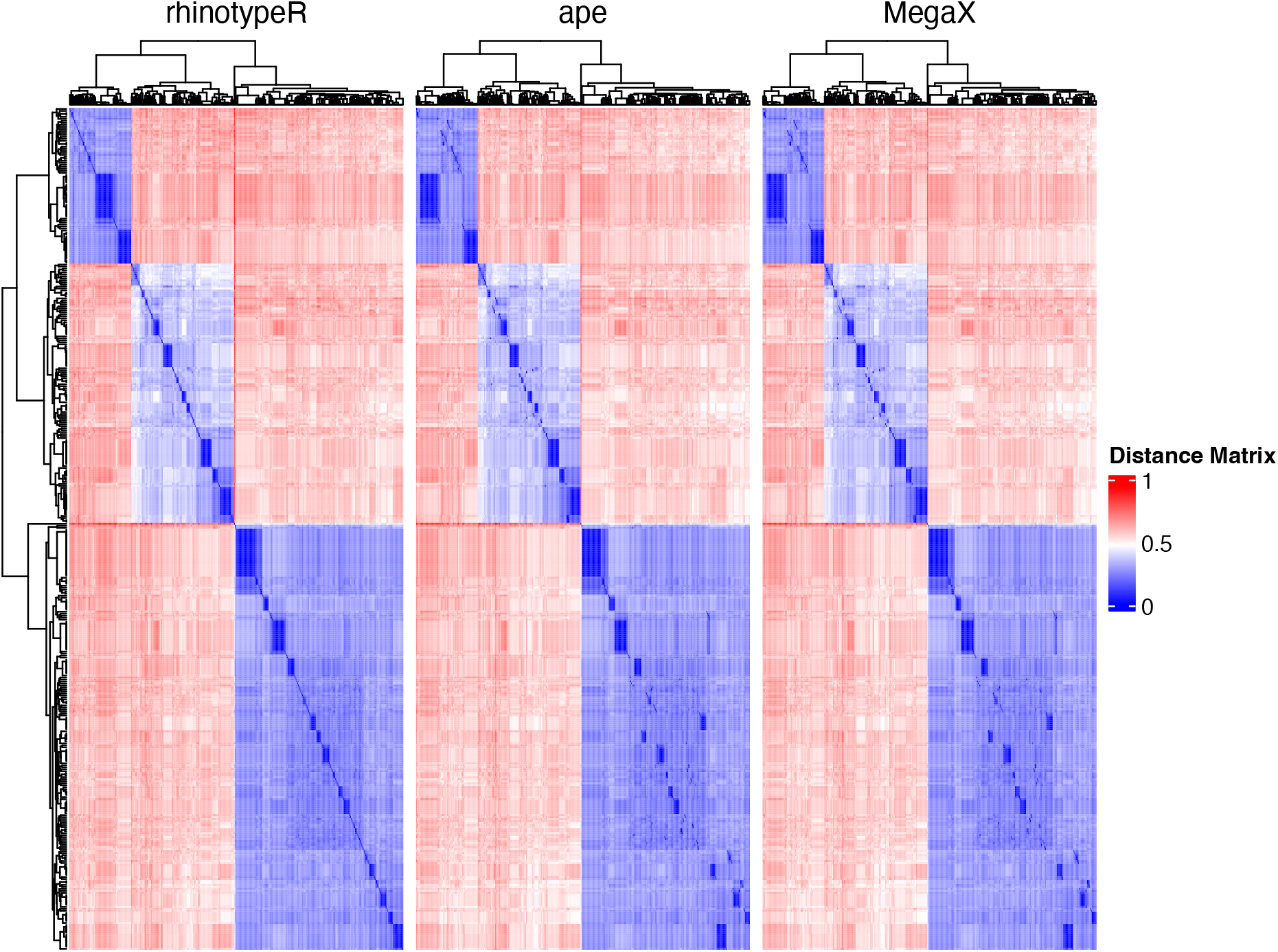
Comparison of the distance matrix generated by rhinotypeR to other commonly used software: the R package ape and Mega X.

The *SNPeek()* function allows users to identify and visualize SNPs relative to a selected reference sequence to detect regions of high genetic variation. Advanced features for examining evolutionary relationships among rhinovirus strains include heatmaps to visualize genetic distances and phylogenetic trees to graphically represent evolutionary connections. Together, these functionalities make rhinotypeR a comprehensive package for rhinovirus genotyping. It offers classification and a suite of analytical and visual tools to aid in interpreting and understanding rhinovirus genetic data.

### Comparative Analysis with Existing Tools

Compared to other tools like RV-Typer^33^, which is no longer accessible, and the rhinovirus database(RVdb) ^34^, which uses the VP1 region and can only process one sequence at a time, rhinotypeR offers a more comprehensive and streamlined approach.

To validate rhinotypeR’s fidelity, we conducted comparative analyses with two established tools: the ape R package^31^ and MegaX^30^, which are commonly used to calculate pairwise genetic distances. Our package demonstrated high congruence with these tools, as highlighted in **Figure 2**. It achieved a correlation coefficient of 0.998 with the distance matrices generated by either tool.

Unlike general-purpose bioinformatics tools that require additional steps to genotype rhinovirus sequences, rhinotypeR automates this process. It collates the prototype strains, calculates the genetic distances, and classifies sequences into genotypes by comparing them against the curated database of prototype strains. This functionality enhances accuracy and significantly reduces the time and effort in rhinovirus genotyping, ensuring reproducibility and reliable results.

RhinotypeR efficiently handles typical datasets with multiple sequences. The tool’s ability to process and analyze extensive data with less manual intervention marks a significant improvement over previous methods constrained by dataset size or required sequential processing of individual sequences.

## 4. Discussion

RhinotypeR represents a significant advancement in viral genomics, offering a streamlined tool specifically designed for rhinovirus genotyping using the VP4/2 region. Filling a critical gap in current bioinformatics resources, rhinotypeR provides a robust, user-friendly solution that facilitates the genotyping process, a foundational step in understanding viral epidemiology and evolution. By automating genetic comparisons, genotype assignment, and visualization, rhinotypeR enables comprehensive exploration of rhinovirus diversity and evolutionary patterns, tailored to the unique challenges of rhinovirus research.

Compared with existing tools, which often lack accessibility or struggle with large datasets, rhinotypeR brings enhanced functionality, allowing researchers to efficiently process extensive rhinovirus data with accuracy. Its ability to generate precise genetic distances and classify sequences against a curated prototype database makes it indispensable for large-scale data analysis. RhinotypeR also minimizes manual data handling, substantially reducing processing time while maintaining a high correlation with established software such as MegaX^30^ and ape^31^.

The intuitive graphical outputs provided by rhinotypeR allow for straightforward visualization and interpretation of complex genetic relationships, enabling researchers to immediately analyze data and contextualize broader evolutionary trends. This capability is especially valuable for rhinovirus surveillance, as it aids in understanding transmission dynamics and could contribute to more informed public health strategies. Given the increasing demand for rapid response tools in infectious disease genomics, rhinotypeR is a timely addition, supporting viral surveillance and public health preparedness by accelerating the analysis of circulating strains.

## 5. Conclusion

RhinotypeR addresses a vital need in rhinovirus research by providing an open-source, efficient, and dedicated genotyping tool. By facilitating accurate, streamlined genotyping and visualization, rhinotypeR saves researchers time and reduces potential errors, paving the way for deeper insights into the genetic diversity and evolutionary dynamics of rhinoviruses. Its development marks a pivotal step forward in viral genomics, enhancing both scientific inquiry and public health response capabilities. Future updates may expand rhinotypeR’s functionality to include newly identified genotypes, additional evolutionary models, or genotyping using the VP1 region.

## Availability and Requirements

*Project home page*: https://bioconductor.org/packages/release/bioc/html/rhinotypeR.html

*Programming language*: R version 4.4+ and BiocManager 3.20+

*License*: MIT

## Acknowledgements

The authors thank the National Institutes of Health (NIH) Office of Data Science Strategy (ODSS) for their immense support before and during the April 2024 Omics codeathon organized in collaboration with the African Society for Bioinformatics and Computational Biology (ASBCB).

## Author Contributions

MML conceived the original idea. MML, WMR, WG, RN developed the pipeline for the project, MML, WG, WMR, RN performed the bioinformatic analysis of the case study data, and drafted the manuscript. MML, WMR contributed in writing, reviewing and editing the manuscript. OIA’s role was in administrating and supervising the bioinformatics analysis in the project. OIA also provided the resources to facilitate and complete the study and provided guidance. OIA edited and reviewed the manuscript and provided critical feedback that helped shape the final version. All authors read and approved the final manuscript.

## Declarations

## Ethics approval and consent to participate

Not applicable.

## Consent for publication

Not applicable.

## Supplementary Material

**Supplementary Table 1**. GenBank Accessions and respective genotypes of prototype sequences incorporated to rhinotypeR

